# A conserved region T-cell vaccine for Sarbecoviruses

**DOI:** 10.1101/2025.05.20.654373

**Authors:** Marta Escarra-Senmarti, Jacqueline Brockhurst, Amanda R Maxwell, Taylor Godwin, Tianle Zhang, Yuzheng Chen, Peter K Um, Ludmila Danilova, Wen-Chi Hsu, Cheng Ting Lin, Tomáš Hanke, William R Bishai, Andrew Pekosz, Cory Brayton, Bette Korber, Maximillian Rosario

## Abstract

The rapid development of vaccines was a critical part of the global response to the COVID-19 pandemic. SARS-CoV-2 (a Sarbecovirus and member of the Betacoronavirus genus responsible for the pandemic) virus was first detected in Wuhan, China in late 2019. Effective mRNA vaccines based on the viral Spike protein were designed from the earliest isolates and available by December of 2020. SARS-CoV-2 has continued to evolve in the human population, accruing neutralizing antibody resistance mutations that have necessitated updating the vaccine periodically to better match contemporary variants. Neutralizing antibody cross-reactivity is generally very limited among the diverse members of the betacoronavirus genus that are of clinical importance in people. Here, we present an alternative vaccine strategy based on eliciting T-cell responses targeting four highly conserved regions shared across the betacoronavirus proteomes. We hypothesized that cross-reactive responses to these regions could temper disease severity. Focusing immune responses on highly conserved epitopes could be beneficial as SARS-CoV-2 continues to evolve, or if a novel betacoronavirus should enter the human population. Vaccination with these highly conserved regions induced robust T-cell responses in mice and rhesus macaques. Vaccinated hamsters were significantly protected against weight loss and lung inflammation after challenge with the SARS-CoV-2 Omicron variant. After a SARS-CoV-2 Delta challenge in rhesus macaques, 3 out of 4 animals in the control group had infectious virus in their bronchoalveolar lavage samples, while the 4 animals in the vaccinated group did not.

## Introduction

Coronaviruses can be divided into two subfamilies (*Coronavirinae* and *Torovirinae*). Coronavirinae are further divided into α, β, γ, and δ genera. It is the β coronaviruses that have given rise to recent outbreaks causing severe respiratory disease among human populations. A subfamily of β coronavirus, Sarbecovirus, is responsible for the COVID-19 pandemic. In late 2019, SARS-CoV-2 crossed the species barrier into humans resulting in a global pandemic causing over 7 million deaths and over 777 million documented infections as of March 2025 (https//:covid19.who.int). During the first year of the pandemic, human populations began to develop immunity through person-to-person spread^1^. This immunity was augmented by effective vaccination campaigns and by early 2023, the US population had reached over 95% seroprevalence^2^. Vaccines have been shown to be beneficial in terms of reducing infections and limiting severe disease and death as well as reducing rates of long COVID^3,4,5^. However, viruses capable of evading neutralizing antibodies^6,7^, whether elicited by prior infection or vaccination, are continuously evolving^8,9,10,11^, necessitating viral variant surveillance and periodic vaccine updates to improve antibody recognition of circulating forms^8,10^. SARS-CoV-2 has also demonstrated that global transitions to a novel variant with new immune resistance and transmission characteristics can be very rapid and take just a few months. On two occasions these shifts were manifested by viruses carrying very distinctive Spike proteins. The first was the rapid transition to the Omicron BA.1/BA.2 variants in the winter of 2021/2022^12,13^, and the second a rapid transition to the BA.2.86/JN.1variants in 2023^14,15^. Each of these transitions involved a new variant that carried more than 30 mutations in Spike protein relative to the variants that were circulating prior to their introduction. Such highly distinctive viral variants may have originated in individuals shedding infectious virus for prolonged periods of time or from unsampled geographic regions. Thus, SARS-CoV-2 has demonstrated the capacity to make unexpected shifts in its evolutionary trajectory to new fitter and immunologically more distinctive viral forms and such transitions can be swift.

To curtail the pandemic, vaccines that maintain and enhance neutralizing responses against currently circulating virus is of great interest^16^. In that effort, vaccination strategies have generally targeted the spike protein. Spike is highly variable compared with other viral proteins both within SARS-CoV-2^17,18^ and among Sarbecoviruses (Fig. 1). The SARS-CoV-2 Spike protein is not only the primary target for neutralizing antibodies, but it is also highly immunogenic in terms of its capacity to induce both CD4^+^ and CD8^+^ T-cell responses^19^. SARS-CoV-2 Spike vaccines induce T-cell responses that to date have maintained much of their cross-reactive potential even as new antibody-resistant variants have sequentially emerged^20^. SARS-CoV-2, however, continues to evolve, and a vaccine that carries highly conserved but otherwise immunologically sub-dominant regions within the viral proteome may have the potential to redirect immune responses towards epitopes that would be stable and relevant against newly arising viral variants. More importantly, the most conserved regions of the Sarbecovirus proteome have the capacity to elicit T-cell responses with the potential for cross-reactivity against other clinically relevant Sarbecoviruses as well as other as yet unknown coronaviruses. Thus, this vaccine strategy could provide a safety net for the uncertainty of future zoonotic events and contribute to pandemic preparedness, as shifting between hosts is a hallmark of RNA viruses such SARS-CoV^21^ and SARS-CoV-2.

**Figure. 1.**
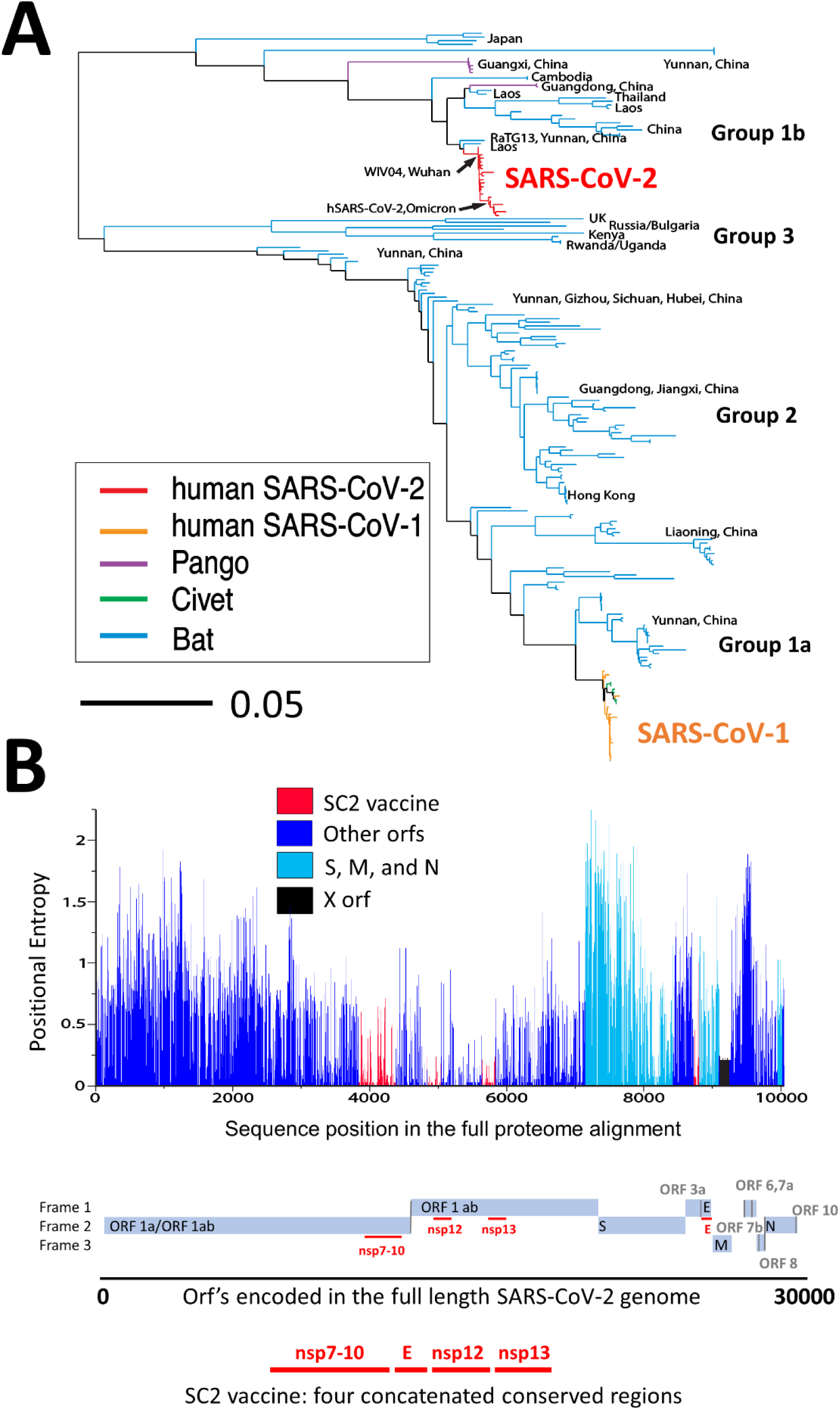
Sarbecovirus diversity. **A.** Phylogenetic tree illustrating the diversity of the Sarbecovirus subfamily. This is a maximum likelihood tree with midpoint rooting, colored to highlight the host species of origin of each of 205 Sarbecoviruses all with full length genomes available; the tree is based on protein sequences using a full proteome alignment and selected to be representative of known diversity. Twenty SARS-CoV-1 and SARS-CoV-2 sequences each were included to be representative of the viral diversity that has been found in humans. Bat populations have been most extensively studied in China, although a small number of samples have been obtained in other parts of Asia, and in Europe and Africa. Geographic origins of subclades are noted in the tree; a tree ordered list of sequence accession numbers, geographic origin, sampling year and host is provided in Supplement Table S1. **B.** Per-site amino acid entropy comparing the conserved regions included in the SC2 vaccine to other regions in the full proteome. The Spike (S), Membrane (M), Non-Structural Proteins (nsp) and Nucleocapsid (N) proteins are highly expressed and tend to be immunodominant for CD8^+^ T cells^30,47^ but they are also highly variable proteins, limiting their cross-reactive potential among Sarbecoviruses outside of SARS-CoV-2. The four red regions are among the most conserved regions in the proteome, and these were the regions which were included in the SC2 vaccine. Aligned underneath the full proteome entropy plot is a graphic showing all of the open reading frame (ORF) coding regions in the SARS-CoV-2 genome, and the relative location and length of the four SC2 fragments; the locations of the ORFS are defined in the SARS-CoV-2 GanBank reference genome (NC_045512.2), and that sequence was also used as the basis for the vaccine sequence. The SC2 vaccine was synthesized as the four concatenated fragments ordered as shown at the bottom: from 5’ to 3’ SC2 sequence (Supp. Fig S1.) includes nsp7-10; Env; nsp12; and nsp13. These four fragments are followed by a Mamu-A*01 epitope, mouse H-2^d^ epitope and Pk tag^48^. 10 pools of 15mer 11-aa overlapping peptides that cover the immunogen were used for immunogenicity testing are illustrated in Supp. Fig. 15A.

While T-cell immunity is unlikely to block infection, it underpins robust adaptive immunity and has proven vital in controlling some viral infections^22,23,24,25,26,27,28^ including coronavirus^29,30,31^. CD4^+^ and CD8^+^ T-cell responses can provide a second defense against diverse variants via cross-reactive and anamnestic protective immunity thus speeding recovery from clinical symptoms^28,32,33,34,35^. Memory T cells targeting the conserved epitopes within SARS-CoV-2 that were likely elicited by other pre-pandemic coronavirus infections were associated with mild symptoms^36^, and a pre-pandemic response to a highly conserved HLA-B*15:01 epitope in Spike was associated with completely asymptomatic SARS-CoV-2 infections^37^. Also of note, memory T cells to conserved epitopes may have resulted in viral clearance among seronegative healthcare workers early in the pandemic^38^. These studies each suggest the potential for protective effects being conferred across diverse Sarbecoviruses.

We hypothesize that the induction of T-cell-mediated immunity through vaccination presents the optimal strategy for targeting conserved regions of the virus. This approach has been evaluated in murine models using adenovirus vectors. Strong cellular immune responses were elicited and, upon challenge, significantly reduced the viral load in the nasal turbinate’s of mouse-adapted SARS-CoV-2^40^. In a pan-Sarbecovirus T-cell vaccine approach, Wee EG et al. ^41^ showed protection in the Golden Syrian Hamster (GSH) model if the vaccine was co-administered with the ChAdOx1 nCoV-19 Spike vaccine at 1/50^th^ the dose. The low fidelity polymerases of RNA viruses^6^ coupled with recombination^43^ provide the underlying viral diversity enabling rapid positive selection. The continuous antigenic drift observed in SARS-CoV-2 at the population level has driven the need to serially update vaccines to enable recognition of contemporary variants^42^. It has also resulted in the loss of sensitivity to most neutralizing antibodies that were used historically as therapeutic reagents, thus requiring sophisticated approaches to enable a breadth of recognition that includes contemporary variants^43^. These issues highlight the potential advantage of incorporating highly conserved regions in vaccine approaches that can help mitigate disease severity.

Here we have employed a conserved proteome strategy targeting internal structures and omitting Spike protein while focusing on hydrophobic regions of SARS-CoV-2 proteins^44,45^. The immunogen we constructed, called SC2, is coded by a codon optimized single reading frame and comprises four concatenated highly conserved regions from nsp7-10, 12, 13 and Env (Fig. 1). By choosing 4 relatively long conserved fragments, we include potential epitopes restricted by many HLAs and minimize the potential for unnatural junctional epitopes.

Pre-clinical testing demonstrated the immunogenicity and protective efficacy of the SC2 immunogen when delivered by combinations of modified vaccinia virus Ankara (MVA.SC2), human adenovirus subtype 5 (HAdV5.SC2), Bacille Calmette-Guérin (BCG.SC2), plasmid DNA (DNA.SC2), and as Synthetic Long Peptides (SLP.SC2). Immunogenicity studies in mice and rhesus macaques (RM) demonstrated specific T-cell response generated by all vector modalities. Mapping HLA class I epitope responses indicated a broad coverage of known Sarbecoviruses. Challenge studies in GSH showed protection with significantly improved weight gain and lung pathology over sham vaccination. In the RM challenge study, the vaccines were shown to be protective by significantly reduced infectious viral loads in the lower respiratory tract.

## Results

*The SC2 Immunogen*. The SC2 immunogen includes four regions of the coronavirus proteome that are highly conserved among known Sarbecoviruses. Sarbecovirus diversity is illustrated in the phylogenetic tree shown in Fig 1A. The immunogen design was based on the first available reference SARS-CoV-2 sequences isolated in Wuhan, China in 2019 at the start of the pandemic (Supp. Fig. S1). While the most highly expressed and immunodominant proteins in SARS CoV-2 are Spike (S), Membrane (M) and Nucleoprotein (N) ^32,46^, these three proteins are also quite variable, which limits their cross-reactive potential between T-cell epitopes found among Sarbecoviruses (Fig. 1B, Supp. Fig. S2), in contrast with the highly conserved regions of the SC2 immunogen. In addition, these conserved regions are nearly invariant among the major lineages responsible for global infections that typify SARS-CoV-2 evolution^42^ (Supp. Fig. S3). This stability is particularly striking when compared to the receptor binding region (RBD) of the Spike protein, that has been under continuing selective pressure from neutralizing antibodies elicited by both infection and vaccination^15,47^. The rate of accrual of amino acid changes among major lineages in the RBD is over 60 times higher than in the SC2 regions (Supp. Fig. S3).

*T cell Immunogenicity in mice*. Immunogen SC2 was expressed from MVA, HAdV5, BCG, plasmid DNA, and also delivered as Synthetic Long Peptides (SLP covering just the conserved regions in nsp12 and nsp13). Correct insertion and expression were assessed by PCR and by adding a monoclonal antibody tag at the C-terminus of the protein (Supp. Fig S4).

Immunogenicity was initially assessed by ELISPOT assay from splenocytes in the BALB/c mouse model (Supp. Fig S4). We tested various regimens of the vectors (Fig 2A and Supp. Fig S5). An MVA.SC2 prime followed by HAdV5.SC2 and then SLP.SC2 boost resulted in the most robust response with the highest average number of pools of peptide (which covered the immunogen, SC2) targeted (7.1 average total) which represented the breath of response Fig 2. On that basis, the MVA-HAdV5-SLP regimen was used moving forward into hamsters and non-human primates. We then tested the polyfunctionality of responses generated from that regimen in BALB/c female mice using FluoroSpot (Mabtech). Re-stimulation with peptides from pools 7-10 resulted in mostly monofunctional T cells that were dominated by IL-2 and TNFα responses (Fig 2B). As a proxy for human responses to the SC2 immunogen, we also vaccinated 37 HLA-A*02:01-transgenic mice (HLA-A*02:01 is one of the most common HLA alleles), 20 males and 17 females, with HAdV5.SC2, pooled the splenocytes of each sex separately, and then mapped responses to a resolution of 15-amino acid-long peptides (Fig. 2C). Vaccination of male and female mice resulted in dominant responses to several epitopes (Fig. 2C). We then vaccinated another cohort of 5 female mice with HAdV5.SC2 and 7 days later assayed for response to re-stimulation with the highest responding peptides from Fig. 2C to confirm 7 dominant CD8^+^ responses Fig 2D. Three of these peptides carried known HLA- A*02:01 epitopes, and the other four contained predicted HLA-A*02:01 epitopes (Supp. Fig. S6). None of these responses were directed toward a junctional epitope, i.e. a peptide that spanned the overlap between two different regions included in the vaccine with a potential to form an unnatural epitope. One peptide was in nsp7, two in nsp8, and four in nsp13, and all seven are highly conserved throughout Sarbecoviruses (Supp. Fig. S6).

**Figure 2.**
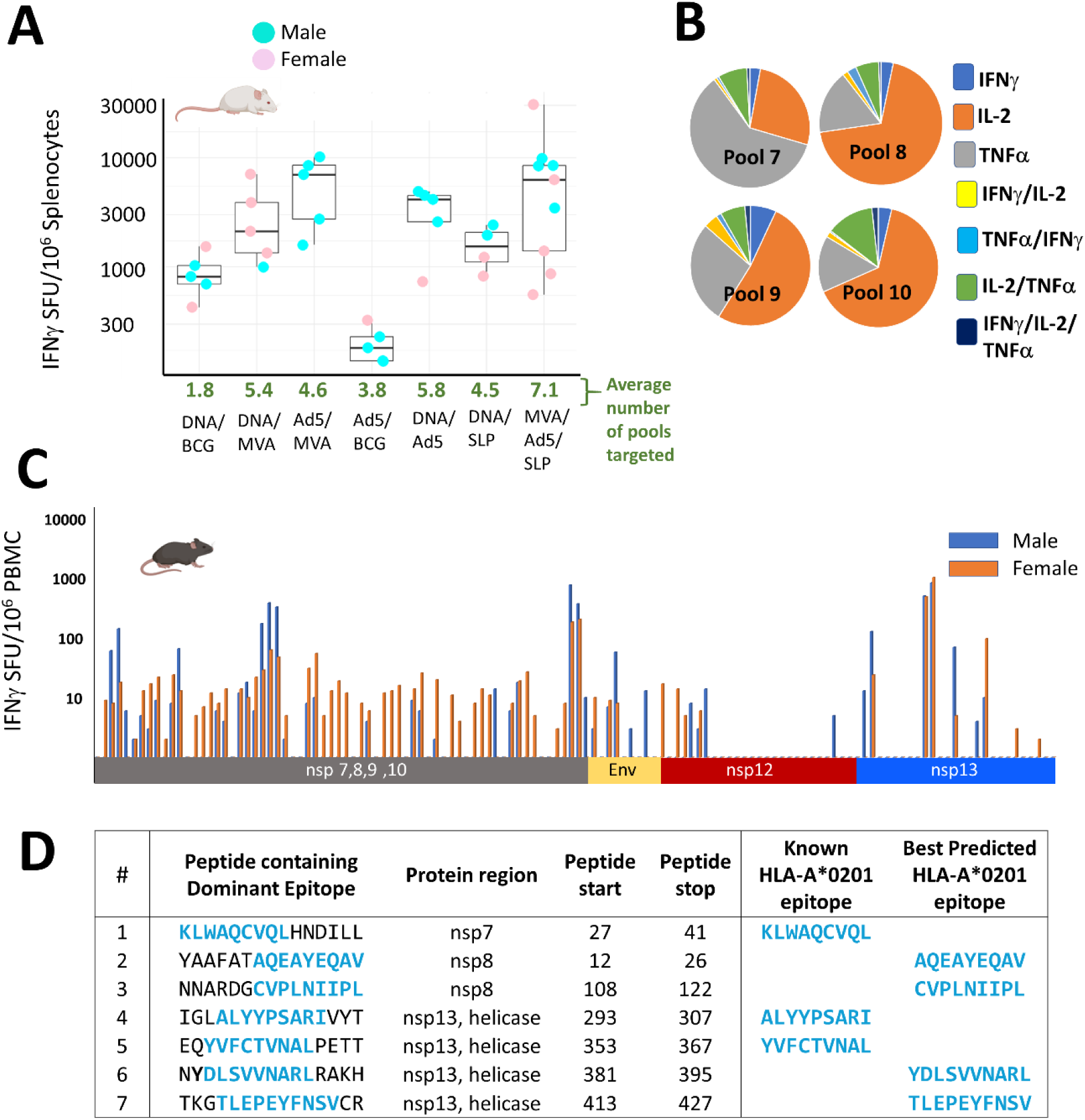
**Vaccine immunogenicity in mice**. **A**. Total IFN-γ responses to regimens of vectors that express SC2 and the order of administration in BALB/c mice. Associated average number of pools of peptides in which responses were noted per group of mice is denoted in green. **B**. FluoroSpot: Average polyfunctional cytokine responses (IFN-γ, IL-2, and TNFα) to re-stimulation with pools of peptides (Supp Fig. S1) from pools 7,8,9, and 10 in five female BALB/c mice. **C**. 20 male and 17 female transgenic homozygous HLA-A*02:01 B6/C57 mice were vaccinated with MVA.SC2 1X10^7^ PFU im. followed by HAdV5.SC2 5X10^7^ PFU im. 21 days later and culled seven days after the boost. PBMC from splenocytes, male or female, were pooled, depleted of CD4^+^ cells (Dynal Beads) and re-stimulated with 10ug/ml of 15mer (11aa overlap) peptides in single wells that covered the immunogen. IFN-γ ELISPOT responses are illustrated (20 SFU/10^6^ PBMC subtracted). **D**. To confirm peak individual CD8^+^ response, a second cohort of five female A*02 mice were vaccinated (with HAdV5.SC2 1 X10^7^ PFU im.) and culled seven days later. PBMC were isolated from spleens and tested against individual peptides with the highest responses seen in C. The animal diagrams were created using Biorender.

*Challenge study in Golden Syrian Hamsters*. We assessed the efficacy of the SC2 vaccines in challenge studies with both Delta B.1.617.2 and Omicron XBB.1.5 subvariants. These variants began their global expantion in the spring of 2021 and in the fall 2022^49,50^, respectively, both after SC2 was designed, thus making them good candidates to test the universal nature of the vaccine. GSH were serially vaccinated with MVA.SC2 10^7^ PFU i.m., HAdV5.SC2 10^8^ PFU i.m., and then SLP.SC2 50 µg/peptide/GSH (divided into two pools administered in the right and left thigh i.m. (Fig 3A). Fourteen GSH were challenged with Omicron XBB.1.5 (1x10^5^ TCID50) and morbidity was determined by percent body mass change from the time of inoculation until 7 days post inoculation (dpi). A significant difference between the two groups was highest at day 5 dpi (p=0.0009) and continued upon recovery at day 7 dpi (p=0.031) (Fig 3B). CT scanning was performed on at 7 dpi to assess inflammation in the affected lung tissue. There was a significant difference between vaccinated and control hamsters as assessed by two blinded radiologists (p=0.00012) (Fig 3C, Supp Fig S7B). Increased whole lung mass has been associated with worse clinical disease^51,52^. Here, we dissected lung lobes and found a significant difference between the masses of the right middle and caudal lobes with trending differences in the right accessory and cranial lobes (Fig 3D). Their left lungs were collected and fixed for histopathology (Supp Fig. S8+S9). Estimates and ranking correlated well with manually outlined areas of H&E involvement, and with Ionized Calcium-Binding Adapter Molecule 1 IBA-1 staining scores (Fig 3C) (Student T-test).

**Figure 3.**
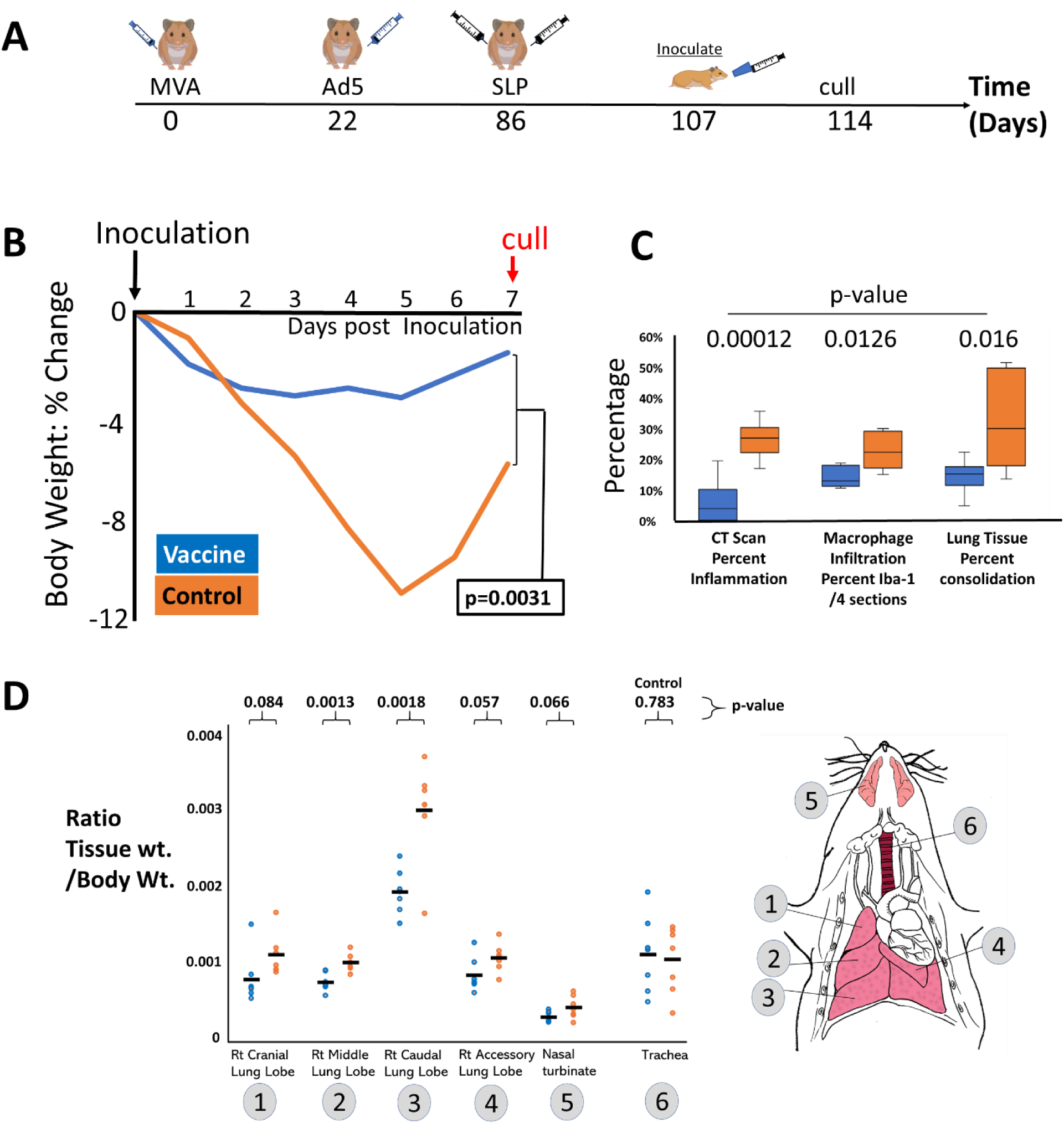
SC2 vaccine efficacy in a Golden Syrian Hamster Challenge Study. Seven female vaccinated (MVA.SC2, HAdV5.SC2, SLP.SC2) and seven sham vacinated (vector/adjuvant only) hamsters were compared. **A**. Schema for vaccination and challenge SARS-Co-V-2 (Omicron). **B**. Morbidity assessed by percent change in body mass from pre-inoculation to 7 dpi. **C**. Percent lung opacity assessed by CT scans, percent left lung macrophage infiltration assessed by IBA1 immunohistochemistry from 4 sections, and percent left lung consolodation at 7 dpi assessed by (H&E). **D**. Lung lobes and turbinate mass at 7 dpi. The animal diagrams were created using Biorender.

Of note, in a challenge study comparing male with female hamsters, post SC2 vaccination with a Delta variant, we saw protective efficacy in the female (p=0.048) but not in the male hamsters (Supplementary figure Supp. Fig S10) based on weight loss after challenge. Others have also observed sex differences in outcomes post coronavirus challenge in the GSH but the mechanisms underlying those differences have not been identified^52,53,54^.

*Challenge study in rhesus macaques*. We assessed the immunogenicity of the most immunogenic regimen of the SC2 vaccines observed in mice (Fig. 2) in the non-human primate model. Four RMs were vaccinated with MVA.SC2, then boosted with HAdV5.SC2 28 days later, followed by SLP.SC2 (divided into four aliquots and administered in the right and left arm and thigh) i.m. 42 days later (Fig. 4A). Polyfunctional CD4^+^ and CD8^+^ T-cell responses to all ten cognate peptide pools were observed (Fig. 4B). These polyfunctional responses to antigen stimulation have been associated with less severe disease in many viral infections^55^ and in convalescent patients polyfunctional memory T cells are associated with less severe disease^19^.

**Figure 4.**
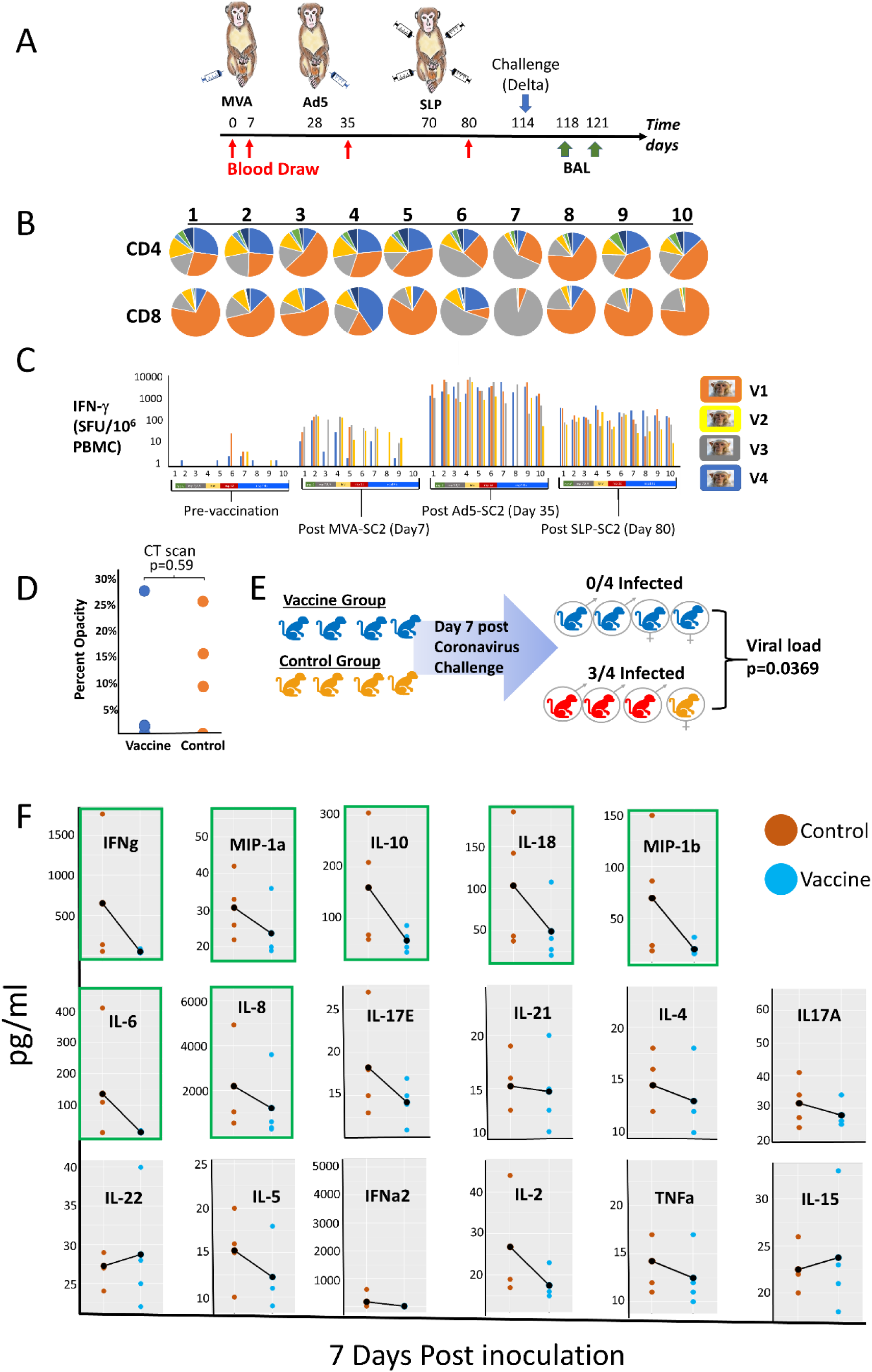
**Immunogenicity and challenge studies in rhesus macaques RM**. **A**. Vaccine schema for eight RM’s. Four RM’s (two males and two females) were vaccinated with MVA.SC2, HAdV5.SC2, and SLP.SC2 and four macaques (three males and one female) received sham vaccines. **B**. Average flourospot value for three assessed cytokines (IFN-γ, IL-2 and TNF-α) at day 17 post SLP-2 vaccination. Cytokine response from PBMC re-stimulated with peptides (15mers with 11aa overlap) covering the immunogen in ten pools (Supplementary Figure 4) illustrates singlet **(IFN-g, IL-2, TNFα),** doublet **(IFN-g/IL-2, IFN-g/TNFa, TNFa/IL-2)** and triplet **(IFN-g/IL-2/TNFa)** cytokine producing cells among CD4^+^ and CD8^+^ subsets. **C**. IFN-γ ELISPOT responses to peptide re-stimulation of PBMC from 4 vaccinated RM’s 7 days prior to vaccination, 7 days post MVA.SC2 vaccination, 7 days post HAdV5.SC2 vaccination, and 10 days post SLP.SC2 vaccination. **D**. Percent opacity as a measure of lung inflammation observed at 7dpi on CT scan. **E**. By day 7 post inoculation, infectious virions from BAL samples were detected in three of four macaques in the control group and zero of four in the vaccine group. **F**. At days 4 and 7dpi post inoculation with Delta (B.1.617.2) variant Luminex (Millipore) multiplex assay was performed on Bronchoalveolar lavage samples from all eight macaques. Cytokines are assessed comparing vaccine with control groups at 7dpi.

The CD8^+^ T-cell IFN-γ release in response to coronavirus antigen stimulation is associated with protection from severe disease^56,57^. Here, we used ELISPOT to assess both the frequencies and breath of T cells to the SC2 immunogen (Fig. 4C and Supp Fig. S11) and also predicted the extent of protection from known viruses based on the recognized epitopes (Supp. Fig. S2).

Post-vaccination with the SC2 vaccine regimen, all four RM’s were bled, and their CD8^+^ PBMCs were isolated to test by IFNγ-ELISPOT assay for responses to peptides spanning the SC2 immunogen (15mers with 11-aa overlap). The assay revealed peak responses in specific regions of SC2 (Supp Fig S11), and a subsequent blood draw (one week later) was used to confirm responses to these individual peptides (considered dominant). We confirmed the dominant responses against nine peptides in RM **V4** (8/9 were highly conserved among Sarbecoviruses), four peptides in RM **V3** (3/4 were highly conserved), and three peptides in RM **V2** (2/3 were highly conserved) and three in RM **V1** (all three were variable) (Supp. Fig. S12).

One of the variable peptides (peptide 3 in nsp9, Fig. S10) was targeted in all four animals. This peptide is quite distinctive in SARS-CoV-2 and closely related viruses relative to other Sarbecoviruses. Despite this, three of the four animals elicited multiple CD8^+^ T-cell responses that were predicted to be highly cross-reactive across all Sarbecoviruses. We also show that all macaques were infected by the challenge virus evidenced by the development of spike specific antibodies only after inoculation, and that these responses were equivalent between the vaccinated and control groups (Supp. Fig S13).

Post a SARS-CoV-2 Delta (B.1.617.2) challenge, a CT scan demonstrating lung opacity at 7 dpi did not show a difference between vaccine and control groups (Fig 4D, Supp Fig S7), however, the macaque with 27.5% opacity (**V4**) in the vaccine group was born with a reduced lung volume. Imaging showed significantly lower lung volume (13.7 cm^3^) vs predicted lung volume (21.4 cm^3^) (OsiriX) for his age (more than 1.96 standard deviations below the mean). This pre-existing condition may have biased the CT scan against the vaccine group because the other three RMs in the vaccine group had lungs clear of visible congestion or infiltrate. Next, we assessed cytokine levels from bronchoalveolar lavage (BAL) samples taken at 7 dpi (Fig. 4F, Supp Fig S14). No individual cytokine was statistically significantly different between the vaccine and control groups (Student -t test). Interestingly, cytokines associated with increased disease severity IL-6, IL-8, IL-10 IL-18, MIP-1α, MIP-1*β,* and IFN-γ^58,59,60^ trended higher in the control group (Fig. 4F). Also, similar to our results, among intensive-care-unit vs ward patients, lower IL-6 and IL-8 from BAL was associated with a better outcome and lower viral loads as assessed by PCR^61,62^.

Clinical signs of illness including reduced appetite, a hunched posture, pale appearance and dehydration were observed in RM’s infected with SARS-CoV-2 isolate nCoV-WA1-2020^63^. We observed few changes in any of the RMs in either the control or vaccine group, and no changes in body weight or temperature (Supp. Table S2). However, some gastrointestinal symptoms were observed in three of the four control RM but not in the vaccinated group. There were significant differences in viral load in the BAL between the vaccinated and control group, with no infectious virus detected 7 dpi in the vaccinated animals while three of four of the animals in the control group had detectable infectious virus (Fig. 4E), suggesting that vaccination with SC2 resulted in more rapid clearing of the lungs post infection.

The durability of T-cell responses in the two groups was assessed in blood collected at more than 200 days post inoculation. An ELISPOT assay comparing total T-cell response to nine pools of peptides that cover the Spike protein showed no difference between the vaccine and control groups. Therefore, the SC2 vaccines did not interfere with the T-cell response to Spike epitopes generated by infection. However, responses to conserved regions of the virus were muted in the control group, detected in only two RMs at less than 100 SFU/10^6^ PBMC (Supp Fig.S15B) while all four SC2 vaccinated RM elicited robust and broad responses to the immunogen (Supp Fig. S15B).

## Discussion

In this work, we show that a T-cell vaccine can elicit protective immune responses against SARS-CoV-2 in two pre-clinical animal models, GSH and macaques. The vaccines expressing immunogen SC2 target conserved regions of Sarbecoviruses, and offer substantial coverage while avoiding the Spike protein (which is under intense selection pressure by the humoral immune system). We show that the immunogen can be presented to the immune system from multiple vectors, each of which induces cellular immune responses in the BALB/c mouse.

Further, vaccine response could be greatly augmented when administered in a heterologous prime-boost regimen. To simulate human vaccination more closely, we show that broad and robust immunity was elicited in HLA-A*02:01-transgenic male and female mice, and identified a number of highly conserved epitopes that could also be recognized in people that carry HLA- A*02:01 (among the most common HLA alleles).

Challenge studies in the GSH provided exceptional protection against sickness from the Omicron variant as evidenced by CT scan, lung pathology, organ specific weights and total body weight loss studies. In the RM model, a heterologous prime-boost regimen of MVA.SC2, HAdV5.SC2 and SLP.SC2 resulted in robust broad polyfunctional cell-mediated responses.

Peptide mapping showed that each animal generated dominant CD8^+^ T cell responses to between 3 and 12 epitopes within the SC2 immunogen, and that most of the epitopes targeted would be expected to be highly conserved across Sarbecoviruses. BAL assay for infectious virus seven days post challenge indicated that no RM receiving the vaccines was producing infectious virions contrasting the 75% of animals in the control group. Luminex studies confirmed a trend in vaccinated animals’ cytokine response that was similar to human responses that are predictive of a positive outcome. RM induced antibodies to the spike region of the virus were observed after inoculation, but none were observed post-vaccination.

Therefore, protection is attributed to the T cells. Further analysis showed similar antibody and cell-mediated responses to the Spike protein in both vaccine and control groups indicating that SC2 is unlikely to interfere with infection and Spike protein vaccine responses. In addition, similar to the people ^64^, we saw few conserved region CD8^+^ T-cell responses post-infection in the control macaque cohort. The vaccinated RMs, however, retained conserved SARS-specific epitope responses for at least 207 days, the last time point measured.

T-cell immunodominant responses can be a critical outcome of vaccination and associated with mild disease^21,65^. Uniquely, we were able to map CD8^+^ T-cell responses across the entire vaccine immunogen and then confirmed immunodominant epitopes with a second blood draw. This information critically supports the idea of the potential for conserved region T-cell vaccines to enable a pan-coronavirus vaccine that can target a diverse range of coronavirus forms found in nature. In this study we have shown that by incorporating these highly conserved regions of the proteome into a vaccine we can focus the immune response on epitopes that are shared across Sarbecoviruses, and that these responses are beneficial in terms of disease outcomes in animal models. However, to extend the response breadth further and to provide extended vaccine coverage throughout the highly diverse coronavirus family, vaccine cocktails spanning this same conserved region but enabling targeting of relevant members of the coronavirus family could be tested. “Epigraphs” ^66^ provide one such vaccine design strategy. The benefit of using conserved region Epigraphs for enabling breadth of vaccine response for other highly diverse viral pathogens, including filoviruses^67^ and influenza^68,69,70^ has been demonstrated in animal models.

Supplementary Figure S3 shows an alignment of 5 historically major variants of SARS-CoV-2 relative to the Wuhan ancestral strain, highlighting the mutations in the spike protein verses the protein regions targeted by SC2. There were 91 amino acid mutations observed in the Spike protein among the founders of these 5 key linages, but only 3 in the SC2 targeted region, highlighting the stability of this region during the pandemic. Moreover, when comparing immunodominant responses induced by SC2 against the full diversity of Sarbecoviruses, there exists a scarcity of mutations that can evade the vaccine elicited responses. This is further echoed in the hypothetical HLA-A*02:01 response seen in transgenic mice where seven immunodominant CD8+ T cell responses were observed (Supp Fig. S6). Note that CD8^+^ T cell responses can often tolerate some diversity^71^, consequently even variants that carry a mutation in an epitope may still be targeted by these vaccine-induced T cells^72^.

NHP models have provided valuable insight into COVID-19 pathogenesis. These models are clinically relevant to humans in terms of the outbred genetics, similar B and T cell selection, and recapitulation of human SARS disease^62,73^. They include rhesus macaques (*Macaca mulatta*), cynomolgus macaques (*Macaca fascicularis*), and African green monkeys (*Cercopithecus aethiops*)^71^. RM’s are commonly used, and early COVID-19 vaccine studies proved efficacious with this model. Specifically, the Pfizer-BioNTech vaccine was tested in RMs and shown to protect from infectious viral challenge together with reduced detection of viral RNA, and this data helped advance the vaccine to human clinical trials^75^. Similarly, AstraZeneca’s ChAdOx1nCo19 (an Adenoviral vectored vaccine) resulted in reduced viral load from Bronchoalveolar lavage samples which also directly led to human clinical trials^76^. Our results with a T cell vaccine have shown similar protection. Furthermore, SC2 was designed in 2020, prior to the emergence of the Delta and Omicron variants^77,78^. The virus has remained nearly invariant in the conserved region we have focused on, while Spike evolved extensively under neutralizing antibody selection and so required vigilant monitoring^79^. SARS-CoV-2 is continuing to adapt and acquire mutations that confer antibody resistance mutations or that are advantageous to growth^43,80,81^. This underscores the potential value of a conserved vaccine design, as well as the relevance of its potential as a pan-Sarbecovirus vaccine.

GSH anatomy and histopathology have made it a useful model^82^. The GSH recapitulates human SARS disease in terms of replication at high titers in the respiratory tract associated with reproducible lung pathology^83^ as well as resulting in human vaccine clinical trials using a variety of delivery strategies^84^. Here, we show strong protection against Delta and Omicron variants for female, but not for male hamsters. Sex differences may present a barrier to detect protection afforded by T cell vaccination^52^ thus requiring RM’s going forward. Or possibly, this finding offers an opportunity to study clinically relevant sexual dimorphisms in response to T cell vaccines.

An ever-increasing group of patients are receiving B-cell depleting agents for lymphoproliferative diseases and other clinical indications, including rituximab, ocrelizumab, and ofatumumab. Unfortunately, these drugs can be consequential for coronavirus vaccination strategies. For example, preliminary studies have shown that those receiving Rituximab had T cell activity similar to healthy controls when given a COVID-19 vaccine, but they did not produce protective levels of anti-spike IgG and IgA antibodies^85^. Thus, T cell inducing vaccines may have a niche in patients taking anti-CD20 medication.

Amidst the tumult of vaccine hesitancy, the SARS-CoV-2 pandemic continues^86^. The virus continues to mutate at a significant rate especially within the spike protein, including the RBD domain^9^ which is heavily targeted by antibody induced immunity. Hospitalizations and deaths have declined as we have developed immunity at the population level^87^. But, antigenic drift continues, and historically dramatically distinct viruses like Omicron’s and JN.1 have rapidly swept the globe, demonstrating the need for continued vigilance. We believe that a conserved T cell vaccine design buffers against immune evasion generated by responses to infection or previous vaccination^88^. With the emergence of the Omicron variants, it is becoming clear that T cell responses can be maintained while humoral efficacy is lost^89^. Currently, there are several candidate ‘Pan-SARS’ antibody-inducing vaccines at various levels in development^90,91,92,93,94^ including T cell inducing conserved antigens^94^. These T cell approaches have shown promise in hamster challenge studies^41,94,95^ but to our knowledge have not been tested in non-human primates. Although not yet licensed by the FDA, a vaccine focused on T cell mediated immunity as a central feature represents a novel concept in vaccinology and is being intensively studied in numerous infectious diseases^63,96,97,97,98,99^. For SARS-CoV-2, T cell control is well established^29^ and for SARS in general, long lived memory responses are possible^100^. With variant diversity and waning of humoral control^101^, a T cell vaccine strategy remains a viable option as a way to complement antibody-based vaccines for SARS-CoV-2 and curtail disease severity^102^. Such a vaccine could have the potential to provide immune protection should a new SARS variant ever emerge that is readily transmitted in the human population, a continuing threat^103^. Indeed, soon partnering antibody with T cell vaccines could synergize to produce a much more effective regimen. Other emerging respiratory diseases may also be vulnerable to this vaccination strategy.

Limitations: Our macaque cohorts were small consisting of four animals per group. Thus, Luminex studies comparing vaccinees with controls did not reach statistical significance. We have not looked at the TCR usage in vaccine and control macaques. Public clonotypes may cross react with variants providing further protection against diversity^104^. We were not able to sample lung and lymph nodes months post vaccination to determine memory resident T cells that may play a role in long term protection^29^. These studies are required for further study.

## Materials and Methods

### Bioinformatic tools

The maximum likelihood phylogenetic tree shown in Fig. 1A was generated using the tool IQ tree^112^ with simple midpoint rooting Shannon entropy was used to estimate the diversity of each position in an alignment of the Sarbecovirus proteome in Fig. 1B using the Entropy tool at the Los Alamos HIV database (as implemented June 2024, for details see https://www.hiv.lanl.gov/content/sequence/ENTROPY/entropy_readme.html). This quantity assesses the uncertainty in each position, essentially how well you would be able to predict the amino acid in that position in an unknown sample drawn from the same population^113^*SC2 vaccine sequence* The genomic sequence of SC2 was synthesized by Genscript Biotech and cloned from pc3.1(+). It encodes concatemerized SARS-CoV2 structural proteins 6-9, 12, 13 and Env. The sequence was ‘humanized’^114^ and encodes a KOZAK start sequence^115^ and H-2^d^, Mamu-A*01, and Pk-tag (aka SV5) C-terminal tag.

### Preparation of pTH.SC2 DNA Vaccine

SC2 was subcloned into pTH^116^ and cultured in-house. Plasmid DNA for immunization was isolated using an Endo-free Gigaprep (Qiagen) and stored at -80 ^0^C prior to use.

### Preparation of MVA.SC2 Vaccine

Recombinant Vaccinia MVA (Modified Vaccinia Ankara) was generated at the University of Iowa Viral Vector Core (Carver College of Medicine, Iowa City, Iowa) ^117^. A partial fragment of SARS- Cov2 was synthesized and cloned by Genscript into a shuttle plasmid downstream of the modified H5 promoter between two flanking regions containing homology to two adjacent essential genes in MVA and a p11 driven GFP transient marker for selection of recombinants.

Following plasmid transfection and homologous recombination into wildtype MVA, flow cytometry was used to sort GFP containing cells into individual wells on a 96-well plate (performed by Flow Cytometry Core at the University of Iowa). Fresh DF-1 cells were added to each well and the plate was incubated for 3 days at 37°C, 5% CO2. DNA from GFP positive wells was purified and analyzed by PCR to identify cells containing recombinants free of wild type MVA. Two GFP positive recombinants containing the transgene underwent several rounds of plaque selection until a GFP negative recombinant was identified. A single recombinant was amplified by infecting DF-1 cells through several passages on an increasing number of 150mm cell culture plates and then purified by sucrose cushion. The final recombinant virus was analyzed by PCR for identity and the infectious titer of the virus was determined by plaque assay. 10^7^ PFU MVA.SC2 was vaccinated subcutaneously (sq. left leg 100 µl) for macaques and hamsters and 50 µl i.m. (calf) in mice.

### Preparation of HAdV5.SC2 Vaccine

HAdV5.SC2 was made with pacAd5(9.2-100) sub360 viral backbone. This backbone has a fully deleted E1a protein, a partially deleted E1b protein, and a partially deleted E3 protein to make the virus replication deficient. All of our vector preparations are purified by double CsCl protocol and dialyzed and stored in our A-195 buffer. All preparations are titered on HEK 293 cells using the Clonetech Adeno-X titer kits and also tested for replication competent particles (RCA). The recombinant adenoviruses can revert to wild type during virus production, thus packaging replication competent particles (RCA) ^118,119^. For this reason, each new lot produced at the core is tested for the presence of RCA by immuno-staining. 5x10^8^ PFU HAdV5.SC2 was vaccinated subcutaneously (sq. right leg 100 µl) for macaques and hamsters and 1x10^8^ PFU HAdV5.SC2 was vaccinated subcutaneously (sq. right leg 50ul) for mouse experiments.

### Preparation of SLP.SC2 Vaccine

Twenty-eight 25mer amino acids (overlapping by 11aa) covering the immunogen were synthesized by Genscript Biotech to >95% purity. These Long Synthetic Peptides SLP were vaccinated (0.2 mg/peptide/rhesus macaque and 0.05mg/peptide/GSH) (100 µl) adjuvanted by topical imiquimod and Montanide ISA-51 (Seppic.) emulsification, divided into four equal sub- pools and injected s.q. into four anatomically separate sites (right arm, left arm, right leg, left leg) in the RM model. For GSH vaccinations (0.1 mg/peptide) peptides were combined into one pool and vaccinated sq. (200 µl) with Montanide ISA-51. For mice vaccinations 0.1mg/peptide adjuvanted with Addavax (Invivogen) was injected as two pools 50ul left calf and 50ul right calf.

### Preparation of BCG.SC2 Vaccine

*Mycobacterium bovis* (*M. bovis*) Bacillus Calmette-Guérin (BCG) Pasteur (BCG Pasteur WT) originated as a tech transfer between the Institute Pasteur and the Trudeau Institute in 1967. BCG Pasteur was then transferred to Johns Hopkins School of Medicine. rBCG.SC2-KmR was generated by electroporation of episomal plasmid pSD5-hsp60-SC2-KmR into electrocompetent BCG Pasteur WT^120^. Electroporated cells were incubated overnight in complete BD BBL^TM^ Middlebrook 7H9 nutrient broth, supplemented with 0.2% Glycerol, 0.05% Tween80, and 10% BD BBL^TM^ Middlebrook OADC Enrichment (7H9) supplemented with 25 ug/mL Kanamycin. After overnight incubation, transformants were plated on BD BBL^TM^ Middlebrook 7H10 Agar solid medium (7H10) with 50 µg/mL Kanamycin and grown at 37°C. Multiple clones were isolated and confirmed by drug selection, PCR, and restriction digest. A single colony was isolated and expanded in 1 mL of complete 7H9 nutrient broth (with 25 µg/mL Kanamycin for rBCG-SC2- KmR). 1 mL of growing culture was inoculated in 10 mL of fresh 7H9 nutrient broth (with 25 µg/mL Kanamycin for rBCG-SC2-KmR) in a 50 mL conical tube and incubated at 37°C at 200 rpm. When the cultures reached an optical density of OD600 = 1.5-2.0, they were sub-cultured in 12 mL of 7H9 media (with 25 µg/mL Kanamycin for rBCG-SC2-KmR) at a starting inoculum of 0.05 and grown to an optical density of OD600 of 0.80 – 1.10 (4-5 days) at 37°C at 200 rpm. Optical density was assessed in liquid suspension at 600 nm using a BioChrom UltroSpec^TM^ 10 Cell Density Meter. Cultures were pelleted, resuspended in 20 mL ice-cold Gibco 1xPBS (pH 7.4) supplemented with 0.005% Tween80, and pelleted again with the procedure repeated. The final vaccine was adjusted with ice-cold 1xPBS (pH 7.4) supplemented with 0.005% Tween80 until a final OD600 value of 0.10 was reached. For mouse vaccinations 100 µl of 10^6^ CFU was administered 50 µl/calf.

### Invitro SC2 expression

Plasmid transfection. HEK 293A cells were cultured in DMEM supplemented with 10% FBS and 1% penicillin-streptomycin. A suspension of 500 µL containing 0.4 million cells was seeded onto a microscope slide. The slide was incubated at 37 °C for 5 hours to allow cell attachment.

Subsequently, adenovirus infection plasmid transfection was performed using jetOPTIMUS reagent, and the cells were incubated at 37°C overnight. Following incubation, the supernatant was removed, and the cells were fixed with ice-cold methanol for 5 minutes. The cells were then washed three times with PBS and twice with PBS containing 2% FBS (PBS-FBS). They were incubated with PBS-FBS for 15 minutes at room temperature. After removing the supernatant, the cells were incubated with Alexa Fluor 488 anti-V5 Tag Antibody (1:500, Thermofisher) and DAPI for 40 minutes at room temperature in the dark. Post-incubation, the cells were washed three times with PBS-FBS and three times with PBS. Two drops of glycerol were added to the slide, which was then covered with a coverslip. The samples were analyzed using a fluorescence microscope (Nikon Eclipse 80i FITC 96302).

BHK were cultured in MEM supplemented with 10% FBS and 1% penicillin-streptomycin. A suspension of 500 µL containing 0.2 million cells was seeded onto a microscope slide. The slide was incubated overnight at 37 °C to allow cell attachment. Subsequently, cells were infected with MVA.SC2, incubation 37 °C overnight. The staining procedure was performed as described above.

### Animal Studies

All experimental procedures were approved by the Johns Hopkins University Animal Care and Use Committee. Johns Hopkins program of Animal Care and Use is accredited by AAALAC International. Animal care and procedures follow the *Guide for the Care and Use of Laboratory Animals*, 8th ed (NRC 2011), and relevant Animal Welfare Regulations. Procedures involving hazardous agents or material were reviewed and approved by JH EHSP.

Mice. BALB/cJ (RRID:IMSR_JAX:000651 and homozygous B6.Cg-Immp2l^Tg(HLA-A2.1/H2-D)2Enge^/J (RRID:IMSR_JAX:00419), male and female mice received from the Jackson Laboratory.

Mice were anesthetized with isoflurane prior to vaccination. At study end, mice underwent terminal spleen collection under isoflurane anesthesia.

Hamsters. Golden Syrian Hamsters (*Mesocricetus auratus*) were obtained from Inotiv and housed singly. They were transferred to a BSL3 facility one week prior to inoculation. All hamsters (male and female) were 7-8 weeks old at the time of vaccination.

For coronavirus inoculation hamsters were anesthetized via IP ketamine and xylazine (60-80 mg/kg and 4-5 mg/kg respectively). Inoculation was performed by intranasal route with a maximum volume of 75 ul/nare. Control animals received equivalent inoculations with dilution media alone.

At study endpoints hamsters were anesthetized using isoflurane and blood and tissues were collected as previously described^50^.

Rhesus macaques (*Macaca mulatta)*. Eight RM (3 female and 5 male) were enrolled in the study. Seven of the macaques were 3 years old and one was 7 years old. All macaques were of Indian-origin, and seronegative for SARS-CoV-2, measles, simian immunodeficiency virus, simian T-cell leukemia virus, and simian type D retrovirus prior to entering the study. For all procedures, animals were anesthetized with intramuscular ketamine (5–10 mg/kg) or ketamine plus dexmedetomidine (0.025 mg/kg). Macaques were socially housed in pairs for the duration of the study.

For virus inoculations, anesthetized macaques were intratracheally challenged with 2x10^6^ TCID50 SARS-CoV-2 [SARS-CoV-2/USA/MD- HP05660/2021] in 1mL 1X PBS. Blood was collected from the femoral vein into heparinized tubes. Bronchoalveolar lavage (BAL) samples were obtained with a modified catheter mini-BAL method^121^ using three 10mL PBS aliquots (30mL total), and nasopharyngeal samples were collected using two nylon flocked swabs (Puritan Medical Products) placed in 3mL PBS. Cell pellets from respiratory samples were separated from fluid by centrifugation at 800 G, and both aliquots were frozen separately at - 80°C prior to use. Macaques were sedated as previously described for CT imaging.

### SARS-CoV-2 viruses

The Omicron variant SARS-CoV-2/USA/MD-HP40900-PIDYSWHNUB/2022 (XBB.1.5; GISAID EPI_ISL_16026423) and the Delta variant SARS-CoV-2/USA/MD- HP05660/2021 (B..1.617.2; GISAID EPI_ISL_2331507) were grown on Vero-E6-TMPRSS2 cells and infectious virus titer determined as described in^122^.

### Infectious virus determination for SARS-CoV-2

Infectious virus titers in BAL samples were measured by determining the 50% tissue culture infectious dose (TCID50) as described in^122,123^ using Vero-E6-TMPRSS2 cells (Japanese Collection of Research Bioresources Cell Bank catalog number JCRB1819).

### Peptides

Peptides for vaccination were synthesized by Genscript Biotech to 95% purity with no N or C terminal alterations. Peptides for testing pools were synthesized by Genscript Biotech and Aapptec to >90% purity. These peptides were 15mer 11 aa overlapping and all were solubilized in DMSO (Hybri-Max Sigma-Aldridge) and stored at 100ug/ul (-80^0^C) before use^124^.

### Cell Collection and processing

*Mice;* splenocytes isolated by strain (Avantor) and plunger (Falcon) method (ref), then washed with RPMI and PBMC isolated over a Ficol bed (Cytiva), and lymphocytes were resuspended in RPMI 1640 supplemented, with 15% FCS, penicillin/ streptomycin prior to assay.

*Macaque;* Peripheral blood mononuclear cells (PBMC) were isolated using Lymphoprep (Cytiva) cushion centrifugation, washed with RPMI (ThermoFisher), and rested two hours prior to assay^121^. For CD8+ T cell isolation, PBMC were co-cultured with CD4+ (Dynal beads) for 20min at 4^0^C under constant rotation. Cells were then washed with RPMI and resuspended in RPMI 15% FCS containing penicillin/ streptomycin prior to assay.

### The IFN- ELISPOT assay

Mabtech ELISPOT kit procedures were used with Millipore (Millipore-Sigma) MSAIP plates. Cells were plated at 50,000-200,000 cells/well in unicate-triplicate where possible and stimulated with 10 pools of peptides covering the SC2 immunogen at 2 µg/peptide/ml. For single peptide re-stimulations, cells were co-cultured in 10ug/ml peptide. Phytohemagglutinin (Invitrogen) and PMA + ionomycin (Biolegend) were used as the positive control for macaques and mice respectively. Cells were incubated for 36 hours, and a well was considered positive if the average number of spots in the negative control was less than a test well. For CD4^+^ and CD8^+^ T-cell depletions (Supp Fig S16), the procedures and reagents of Dynal magnetic beads were followed^71^.

Spots were quantified in the Immune Monitoring Core at the Johns Hopkins Sidney Kimmel Comprehensive Cancer Center (NCI CCSG P30 CA006973) on a Autoimmun Diagnostika GmbH. For flourospots, 200,000 cells/well were used in Mabtech pre-coated plates.

Fluorescent plates were read using a CTL/Immunospot S2 Universal M2 analyzer (CTL, Cleveland OH) and analyzed using vendor provided (Immunospot 7.0.28.7) software.

### ELISA

For ELISA assays, spike proteins were obtained from BEI resources, NIAID, NIH: NR-55710 AY.1 Lineage (Delta Variant). All Glycoprotein (Stabilized) from SARS-Related Coronavirus 2, with C-Terminal Histidine and Avi Tags, Recombinant from HEK293 Cells.

RM IgG SARS-CoV2 spike specific antibodies were determined using a sandwich assay. Briefly, 100 ng/well of whole spike protein (BEI-Resources NR-55170) in PBS was coated in an ELISA plate (Immulon 2HB) at 4^0^C overnight. The plate was blocked with Casein Blocking buffer (Thermofisher scientific) before serum diluted in blocking buffer was added. The plate was incubated with shaking for 2 hours, washed and detection antibody was added 1:1000 (IgG NHPRR clone 8F1-biotin, Non-human Primate Resource Center: Reagents used in this study was provided by the NIH Nonhuman Primate Reagent Resource (PR-8616, AB_2819299, Anti- primate IgG [8F1]-biotin). After two hours the plate was washed and Ultra Streptavidin-HRP 1:2000 in blocking buffer was added. The plate was washed and 100ul of TMB (Cell Signaling Tech.) substrate was added followed by 100ul stop solution (BioLegend) after 30 minutes. The plates were read at 450nm on a Spectramax iD3 plate reader.

### Cytokine analysis

Cytokine assay from Bronchoalveolar Lavage samples were performed on a Millipore Sigma Luminex platform (MILLIPLEX® Non-Human Primate Cytokine/Chemokine/GF Panel (PRCYASF-40K). Antibodies included IFN-γ, IFNα2, IL-10,IL-2, IL-15, IL-17A, IL-6, IL-8, IL-17E, IL-21, IL-4, IL-18, IL-22, IL-5, MIP-1α, and MIP-1β, and TNFα. The Bioplex 200 platform (Biorad, Hercules CA) was used to determine the concentration of multiple target proteins in the extracted specimens. The methodology used a magnetic bead–based fluorescence sandwich ELISA. Thawed samples were centrifuged at 10,000 RCF for to remove particulates, and then added in triplicate wells to prepared beads, according to the manufacturer’s protocol.

Concentrations were determined using 5 parameter logistic function with 1/*y*^2^ massing fit using Belysa v.1.2 analysis software (Millipore Sigma). The lower limit of quantitation was defined by the software for each analyte.

### Computed Tomography

For the GSH and RM live animals were scanned under biocontainment^125^ in the Center for Infection and Inflammation Imaging Research Core using a nanoScan PET/CT (Mediso, Arlington, VA) for hamsters and CereTom eight-slice CT scanner (NeuroLogica, Denvers, MA) for macaques. Investigators were blinded to vaccine and control grouping. Seven days post infection, SARS-CoV2 infected macaques (n=8) and hamsters (n=14) underwent chest computed tomography (CT). The extent of pulmonary disease on chest CT between the vaccinated and control groups were compared. When present, lung findings consisted of patchy consolidations and ground-glass opacities in a symmetric bilateral distribution. A quantitative approach was used to determine the percentage of lung disease and showed a high concordance with consensus qualitative assessments. CT images were evaluated on lung windowing using OsiriX (Pixmeo, Switzerland) and 3D Slicer (free open-source software package). Both the full lung volume and the diseased volumes of interest (VOIs) were manually segmented. The percentage of diseased lung relative to total lung volume was calculated for each scan. In addition, CT scans were qualitatively and independently reviewed by two thoracic radiologists with 10 and 6 years of post-graduate experience, respectively. Concordance between the quantitative and qualitative measurements was assessed. Imaging interpretations were conducted while blinded to the animals’ vaccination status.

### Histopathology, immunohistochemistry

GSH left lungs were fixed in 10%NBF, trimmed into 3-4mm thick cross sections, four or five per cassette, processed routinely to paraffin, sectioned at 4-5u, stained with hematoxylin and Eosin (H&E), Unless specified otherwise. Glass slides (H&E and IHC) were scanned to SVS formats at 40x on NanoZoomer S210 Digital slide scanner (Hamamatsu USA), and assessed on Concentriq digital pathology platform (Proscia Inc., Philadelphia, PA). H&E stained tissue sections of Hamamatsu 40x scanned digital slides were evaluated and annotated on Concentriq Proscia software by an ACVP boarded veterinary pathologist (CB) in a masked manner, following principles for reproducible tissue scoring. Scoring based on^126,127,128^. Percent affected H&E was estimated in 10% increments of 0.5-1x fields of Hamamatsu scanned slides. Estimates correlated well with manually outlined areas of consolidation. Scoring was practiced/confirmed by calculating % from manually outlined Areas affected/Scorable area (less lumens of large air spaces or vessels). IBA1 immunostained (SVS) digital sides were analyzed in QUpath. Further scoring was based on the most affected section (indicated by annotation in slide scans).

BEI resources: The following reagent was obtained through BEI Resources, NIAID, NIH: Spike Glycoprotein (Stabilized) from SARS-Related Coronavirus 2, Wuhan-Hu-1 with C-Terminal Histidine and Twin-Strep^®^ Tags, Recombinant from CHO Cells, NR-53937. The following reagent was obtained through BEI Resources, NIAID, NIH: Spike Glycoprotein (Stabilized) from SARS-Related Coronavirus 2, AY.1 Lineage (Delta Variant) with C-Terminal Histidine and Avi Tags, Recombinant from HEK293 Cells, NR-55710.

## ACKNOWLEDGEMENTS

We thank the Veterinary staff at Research animal resources, the Center for Infection and Inflammation Imaging Research for CT scans, the Clinical Immunology staff at the Department of Pathology for help with materials and the Immune Monitoring core for ELISPOT analysis. We thank the Hamad and Caturegli labs for experimental advice, HEK-293 cells and use of a Flow cytometer. Viral vectors were provided by the University of Iowa Viral Vector Core. This work was supported by the Bisciotti Foundation, Lya and Harrison Latta Fund for the Advancement of Pathology, the Covid-19 Preclinical Research Discovery Fund at Johns Hopkins, the non-human primate reagent resource center and the Johns Hopkins Center of Excellence in Influenza Research and Response (contract HHS 75N93021C00045). BK was supported by grants from the National Institutes of Health, 1P01AI158571 and the Bill & Melinda Gates Foundation INV-042469.

## AUTHOR CONTRIBUTIONS

Vaccine design: B.K and M.R,; Conceptualization and oversite M.R.; Immunogen production, M.E.S. P.K.U., W.R.B., T.H. and M.R.; mouse husbandry and immune studies, M.E.S. and M.R.; hamster husbandry and challenge studies, J.B., A.P., and M.R.; hamster pathology studies, T.G. and C.B.; macaque husbandry, immune and challenge studies, J.B., A.M., A.P., and M.R.; viral load assays, T.Z., G.Z., and A.P.; Bioinformatics, B.K.,; Statistics, L.D.,; Radiology, W.C.H. and C.T.L.; Writing-original draft B.K. and M.R.,; review and editing, all authors.

